# BMP signaling pathway member expression is enriched in enteric neural progenitors and required for zebrafish enteric nervous system development

**DOI:** 10.1101/2023.09.24.559191

**Authors:** Joshua A. Moore, Arielle S. Noah, Eileen W. Singleton, Rosa A. Uribe

## Abstract

The vertebrate enteric nervous system (ENS) consists of a series of interconnected ganglia within the gastrointestinal (GI) tract, formed during development following the migration of enteric neural crest cells (ENCCs) into the primitive gut tube. Much work has been done to unravel the complex nature of extrinsic and intrinsic factors that regulate processes that direct the migration, proliferation, and differentiation of ENCCs. However, ENS development is a complex process and we still have much to learn regarding the signaling factors that regulate ENCC development. Here in zebrafish, through transcriptomic, *in situ* transcript expression, immunohistochemical analysis, and chemical attenuation, we identified a time-dependent role for bone morphogenetic protein (BMP) in the maintenance of Phox2bb^+^ enteric progenitor numbers and/or time of differentiation of the progenitor pool. In support of our *in silico* transcriptomic analysis, we identified expression of a novel ENS ligand-encoding transcript, *bmp5,* within developmental regions of ENCCs. Through generation of a novel mutant *bmp5^wmr2^,* we identified a functional role for BMP5 in proper GI tract colonization, whereby *phox2bb*^+^ enteric progenitor numbers were reduced. Altogether, this work has identified time-dependent roles for BMP signaling and identification of a novel extrinsic factor, BMP5, that is necessary for proper formation of vertebrate ENS.

## Introduction

The enteric nervous system (ENS) is an intrinsic, autonomous network of neurons and glial cells that span the length of the gastrointestinal (GI) tract and controls its intrinsic functions. This expansive network of enteric neurons in vertebrates rivals the number of neurons found within the spinal cord, estimated to be in in the hundreds of millions (Furness J.B., 2006). In the ENS, neurons and glia are arranged in functional units called ganglia. Numbering in the thousands, layers of enteric ganglia are interconnected and lie sandwiched between the two muscular gut layers, the longitudinal and circular, that regulate peristaltic function. As the largest part of the peripheral nervous system, the ENS consists of a large variety of neuronal subtypes, and electrophysiologically-classified cellular identities working in concert to locally orchestrate control of the GI tract (Rao and Gershon, 2018).

Remarkably, the vertebrate ENS is primarily derived from the neural crest cell (NCC) population. NCCs are a highly migratory, proliferative, and multipotent stem cell population of embryonic stem cells that delaminate from the dorsal neural tube, migrate throughout the embryo, and give rise to dozens of tissues and cellular populations. In jawed vertebrates, NCCs emigrate from the vagal level of the neural tube through numerous microenvironmental niches that serve to orchestrate the migratory, proliferative, and differential potentials of this transient stem cell population (Elworthy et al., 2005; Epstein et al., 1994; Hutchins et al., 2018; Kuo and Erickson, 2011; Le Douarin and Teillet, 1973). The orchestration and formation of the complex ENS network has been studied to understand NCC migratory potential and nature of enteric progenitor populations. To this end, elucidation of the timing of the vagal NCC migratory events is well understood in animal models, such as mouse and chick (Le Douarin and Teillet, 1973; Tang et al., 2021). Vagal NCC populations immigrate into the developing foregut, where they are referred to as enteric neural crest cells (ENCC), and continue along its length to colonize the gut, through the hindgut. ENCCs then differentiate into enteric neurons or glial cells.

In zebrafish, a robust vertebrate model that has become popular for ENS studies (Ganz et al., 2016), once vagal NCCs enter the foregut at ∼ 32 hours post fertilization (hpf) they are known as ENCCs (also known as enteric progenitors). The ENCCs migrate caudally in two chains from the foregut through the hindgut of the zebrafish gut and express the markers *phox2bb* and *sox10* (Elworthy et al., 2005; Harrison et al., 2014; Uribe and Bronner, 2015; Taylor et al., 2016). Colonization of the gut by enteric progenitors can be fully visualized using live cell imaging and confocal microscopy, and by 120 hpf, the zebrafish ENS contains differentiated enteric neurons along its length (Ganz et al., 2016). Despite its relatively simple gut morphology and ENS structure (Ganz et al., 2016; Willms and Foley, 2023), zebrafish share significant functional conservation of morphological, genetic, and gut physiological similarities to humans.

Currently, we lack comprehensive understanding of the multifactorial network of extrinsic and intrinsic factors that orchestrate the specification, differentiation, and proliferative capacity of ENCCs. To date, much research attention across organisms has centered upon a signaling pathway that is necessary for proper ENS colonization, Ret, a receptor tyrosine kinase. Ret responds to the binding of TGF-β signaling factor, glial cell-derived neurotrophic factor (GDNF), and promotes the migration of enteric progenitors (Natarajan et al., 2022). Recent work in zebrafish and mouse demonstrates more complex roles of traditionally studied ENS factors, such as Ret (Baker et al., 2022; Natarajan et al., 2022; Vincent et al., 2023), where it was recently found to be necessary for driving not only ENCC migration but the proliferative capacity and population density of enteric neurons along the gut. While these recent efforts have extended our understanding of how certain previously known pathways, such as Ret, mechanistically regulate ENCC ontogenesis, delineating how other known factors may be involved in specific phases ENS formation have not been as well studied.

One type of protein that is important for ENS development is Bone Morphogenetic Protein (BMP), a class of TGF-β signaling molecules. BMPs are major developmental morphogens that are involved in many aspects of development from invertebrate to vertebrate organisms (Madamanchi et al., 2021; Wang et al., 2014; Webster et al., 2021). Previous work in zebrafish (Huang et al., 2019), humans (Memic et al., 2018; Wu et al., 2014), and other amniote models (Chalazonitis et al., 2008, 2004; Goldstein et al., 2005), suggest that BMPs, outside of the context of neural crest specification, are necessary for proper ENS colonization and specification. Specifically, broad attenuation of BMP signaling has demonstrated the involvement of BMPs in proper intestinal colonization by the ENCC to form the ENS (Fu et al., 2006; Goldstein et al., 2005) however, the specific BMP ligands and timing that could drive this process has yet to be clearly identified. More recently, transcriptomic analysis of human and mouse intestinal tissue predicts the expression of numerous BMPs and their receptors in the developing GI tracts (Memic et al., 2018), thereby suggesting functional roles for BMPs throughout the development of the ENS.

Within this work here, we define the spatiotemporal expression of BMP pathway members, BMP target genes, and resolve a temporal requirement for BMPs during early ENS development in zebrafish. Specifically, we identify a role that the BMP pathway plays in controlling the timing of ENS neuronal differentiation. We also leveraged mutagenesis of a novel ligand-encoding gene, *bmp5*, in order to enhance our understanding of BMP effects on how the ENS is established during early development.

## Results

### BMP receptor and ligand transcripts show distinct transcriptomic expression in enteric progenitors and neurons

To elucidate which BMPs are expressed during zebrafish ENS formation we queried our previously published posterior *sox10:*GFP single-cell transcriptomic datasets (Howard et al., 2021), which encompass 48-50 and 68-70 hpf time points (Fig.1A). 37 BMP signaling pathway members and target genes were identified (Supp. Fig.1) across *sox10:*GFP^+^ neural crest, enteric progenitors, and enteric neuronal populations (Fig.1A). These populations were identified by the expression of major conserved transcriptomic identifiers *sox10*, *phox2bb*, and *elavl3* (Fig.1B) as previously demonstrated (Elworthy et al., 2005; Howard et al., 2021; Martik and Bronner, 2017; Taylor et al., 2016). We identified BMP pathway member transcripts, such as *bmp4* and *bmp5,* encoding BMP ligands (Fig. 1C); and *bmpr1ba, bmpr1aa*, *acvr1l* and *acvr2aa,* encoding BMP Receptors (Fig. 1D) across *sox10*:GFP^+^ neural crest, enteric progenitors, and neurons. These data support the expression of BMP signaling pathway members across the populations of cells that give rise to the ENS and recapitulate broad expression of BMPs across other amniote enteric populations (Memic et al., 2018). The identification of ligand, receptor, and target gene transcripts supports a functional role for BMPs in the development of zebrafish ENS.

**Figure 1.**
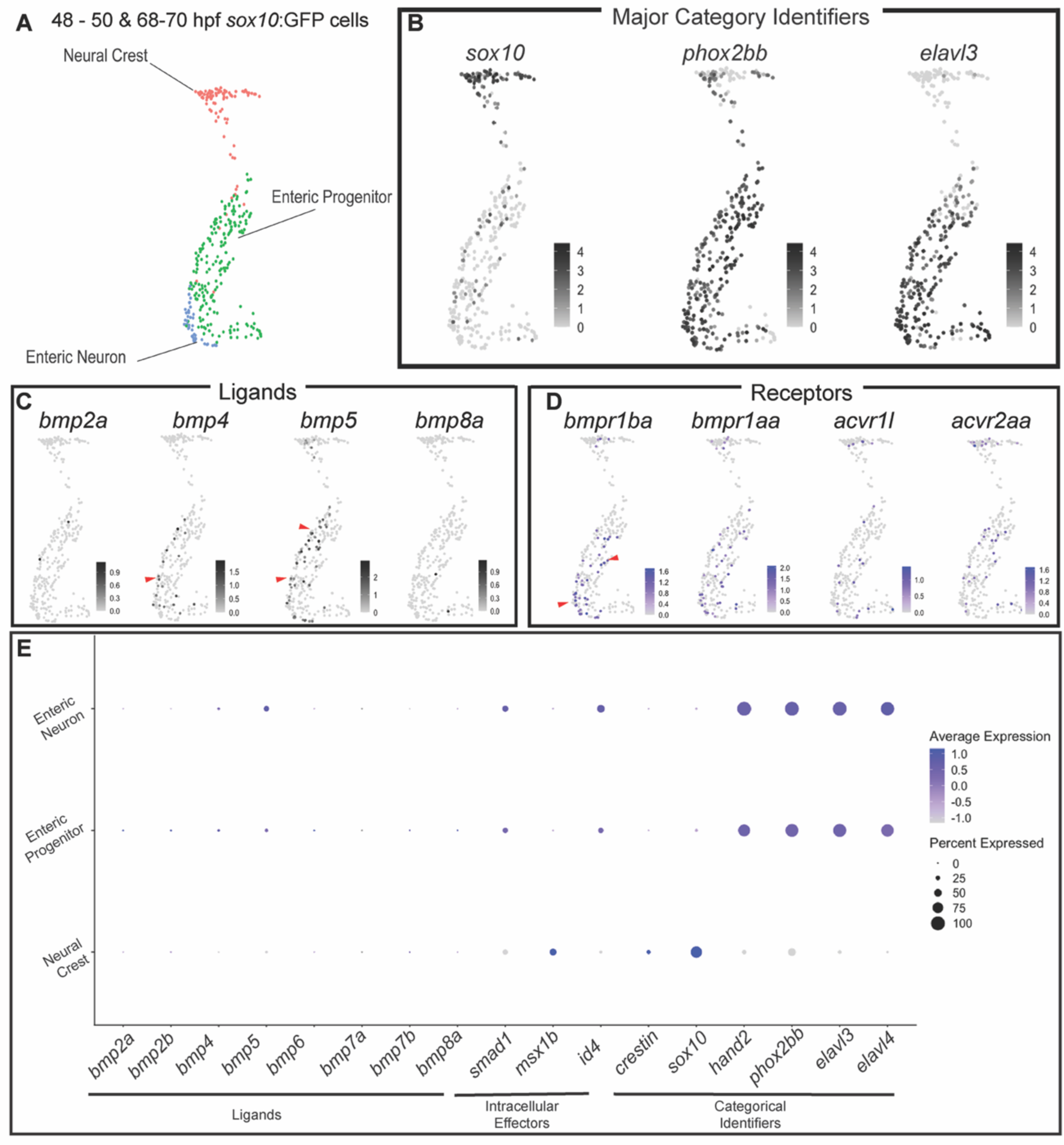
BMP Pathway members are expressed in various *sox10*:GFP^+^ populations, including neural crest and enteric neural progenitors. UMAP analysis of posterior *sox10*:GFP^+^ neural crest, enteric progenitors, and enteric neurons from combined 48-50 and 68-70 hpf time points from a previously published data set (Howard et al. 2021) (A). Feature Plots demonstrate major categorical identifiers of derivatives (B), expression of BMP ligands (C), or receptors (D). Dot plot depicts the expression of BMP ligands, intracellular effectors, and categorical identifiers (E). Deepness of color denotes higher relative expression of genes. Dot size denotes the percentage of the population expressing the gene of interest.

### Target genes of the BMP signaling pathway are expressed in enteric progenitors

Previous studies have identified several downstream genes of BMPs responsible for various aspects of the developmental processes. For example, the family of Muscle Segment Homeobox genes (MSXs) are involved in dorsal-ventral patterning (Esteves et al., 2014; Pomreinke et al., 2017) and neural crest specification (Tríbulo et al., 2003). Along with this, recent works identified the inhibitor of DNA binding family of proteins are involved in migration of cancerous cells (Ke et al., 2018), and are direct targets of BMPs in embryonic stem cells (Hollnagel et al., 1999; Shih et al., 2017). Interestingly, specific members of the ID family of proteins, like ID2a, have known roles during maintenance of the NCC mesenchymal transition (Das and Crump, 2012), and for neurogenic capacity during embryonic retinal development (Uribe and Gross, 2010).

To elucidate whether *id* and *msx* gene family members were expressed in NCC and enteric cells, we queried their expression in our single-cell *sox10*:GFP^+^ datasets. We found *msx* and *id* genes expressed in neural crest, enteric progenitor and enteric neuron populations (Supp. Fig.1; Fig.1C). We then utilized hybridization chain reaction (HCR) to determine spatiotemporal expression patterns of cells that were potentially responsive to BMPs across various time points of relevance (Fig.2A, B). We identified the co-expression of zebrafish-specific neural crest marker *crestin* and BMP target transcript *msx1b* at 24 hpf (Fig.2C-E) and 36 hpf (Fig. 2F-H). When looking at the regions of *msx1b* expression, we identified dorsal expression at 24 hpf (Fig.2C-E, arrowhead) and near the anterior developing foregut at 36 hpf (Fig.2F-H, arrowhead). When looking later we identified co-expression of ENS marker *hand2* (Howard et al., 2021; Reichenbach et al., 2008) and BMP target gene *id2a* at 48 hpf (Fig.2I-K) and 72 hpf (Fig.2L-N). This HCR-assayed time-dependent target gene transcript expression is supported by single-cell transcriptomic analysis across early developmental time points, including 24 hpf (Fig. 2O) (Howard et al., 2021; Lencer et al., 2021). The identification and expression of *msx1b* and *id2a* suggest a spatiotemporal-dependent expression of BMP target genes during both the early and late phases of ENS development.

**Figure 2.**
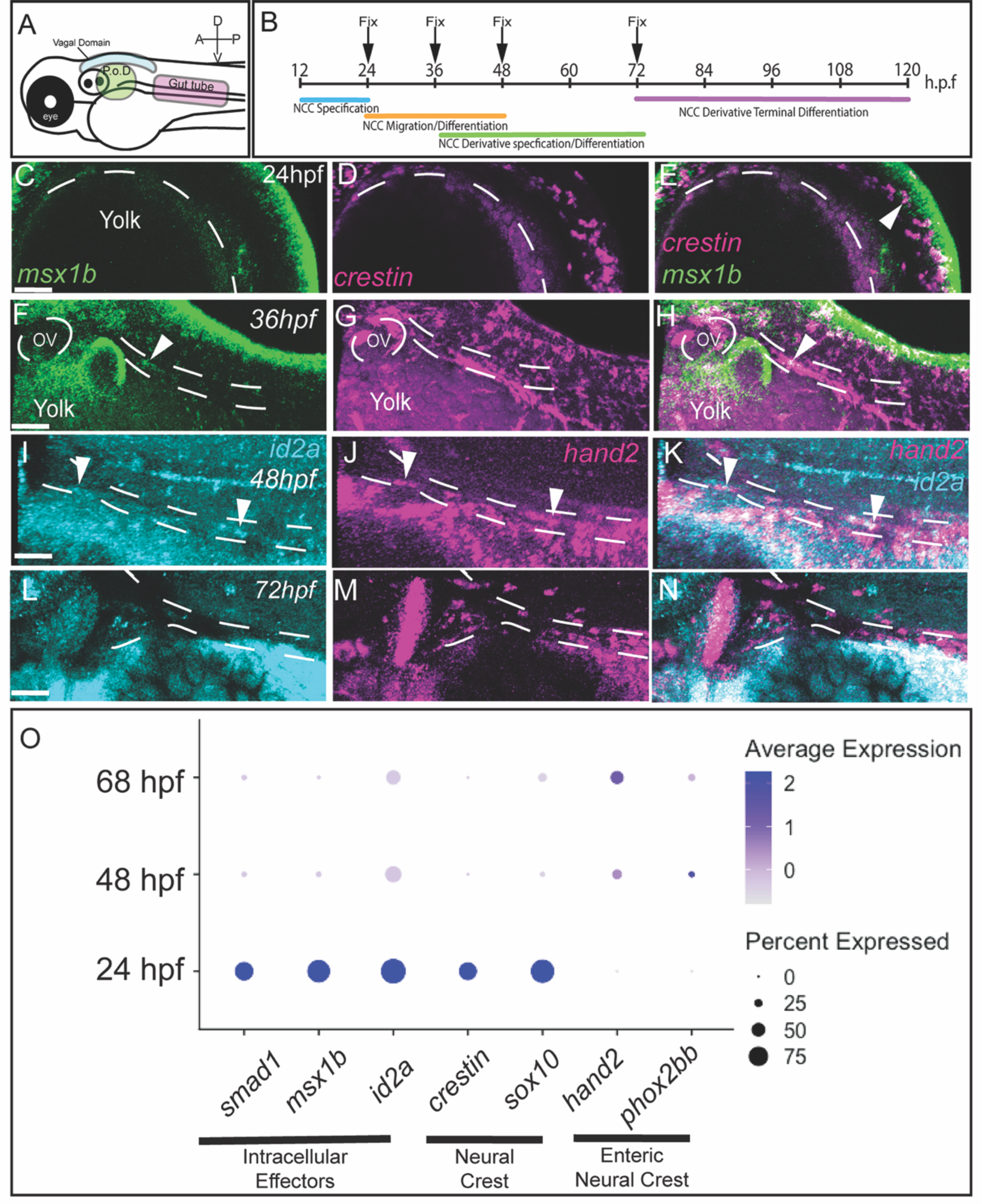
BMP Pathway target genes are expressed in *crestin*^+^ and *hand2*^+^ progenitors. Schematic of regions of interest in the zebrafish embryo (A). HCR analysis of *msx1b* and *crestin* co-expression at 24 hpf (C-E) and 36 hpf (F-H), or *id2a* and *hand2* co-expression at 48 hpf (I-K) and 72 hpf (L-N). White arrows denote regions of co-expression along the dorsal aspect (E), or in the gut tube (F-K). Dot plot demonstrates the expression of select cell identity and BMP pathway categorical identifiers with intracellular effectors across major developmental times (O). Anterior oriented to the left and dorsal to the top, in these images. The scale bar depicts 50 microns in C, F, I, L. P.o.D.: Post-otic Domain.

### BMP-specific R-Smad activation occurs in Phox2b^+^ cells during embryogenesis and enteric nervous system development

Our single-cell transcriptomic and HCR analyses (Fig.1,2) predict BMP signaling activity during development of the zebrafish ENS. To validate the predicted activity of the BMP pathway in NCC and enteric progenitors, we utilized immunohistochemical analysis of activated BMP-specific Smad 1/5/9 (Das and Crump, 2012) and Phox2b (Howard et al., 2022) (Fig.3A,B). Phox2b and phosphorylated R-Smad 1/5/9 (pSMAD) lacked co-expression at 24 hpf (Fig.3C-E’), though BMP signaling showed activation in regions of neural crest migratory paths along the vagal-level neural tube. Co-stain of Phox2b and pSMAD was evident at ∼32-36 hpf in cells along the post-otic domain (P.o.D.) (Fig. 3 F-H’, arrowheads). This pool of cells is thought to be the source of the migratory chain of enteric progenitors that migrate to populate the developing gut (Anderson et al., 2006; Hutchins et al., 2018; Uribe and Bronner, 2015; Young and Newgreen, 2001). Phox2b and pSMAD co-expression was maintained through 48 hpf (Fig.3I-K’), however, co-expression dissipated by 72 hpf along the gut (Fig. 3L-N’). These temporal changes in co-expression suggested the potential of time-specific functional roles for pSMAD-mediated BMP signaling in enteric progenitors. Of particular interest is insight into how BMP could affect ENS development, perhaps by the timing of BMP activation.

**Figure 3.**
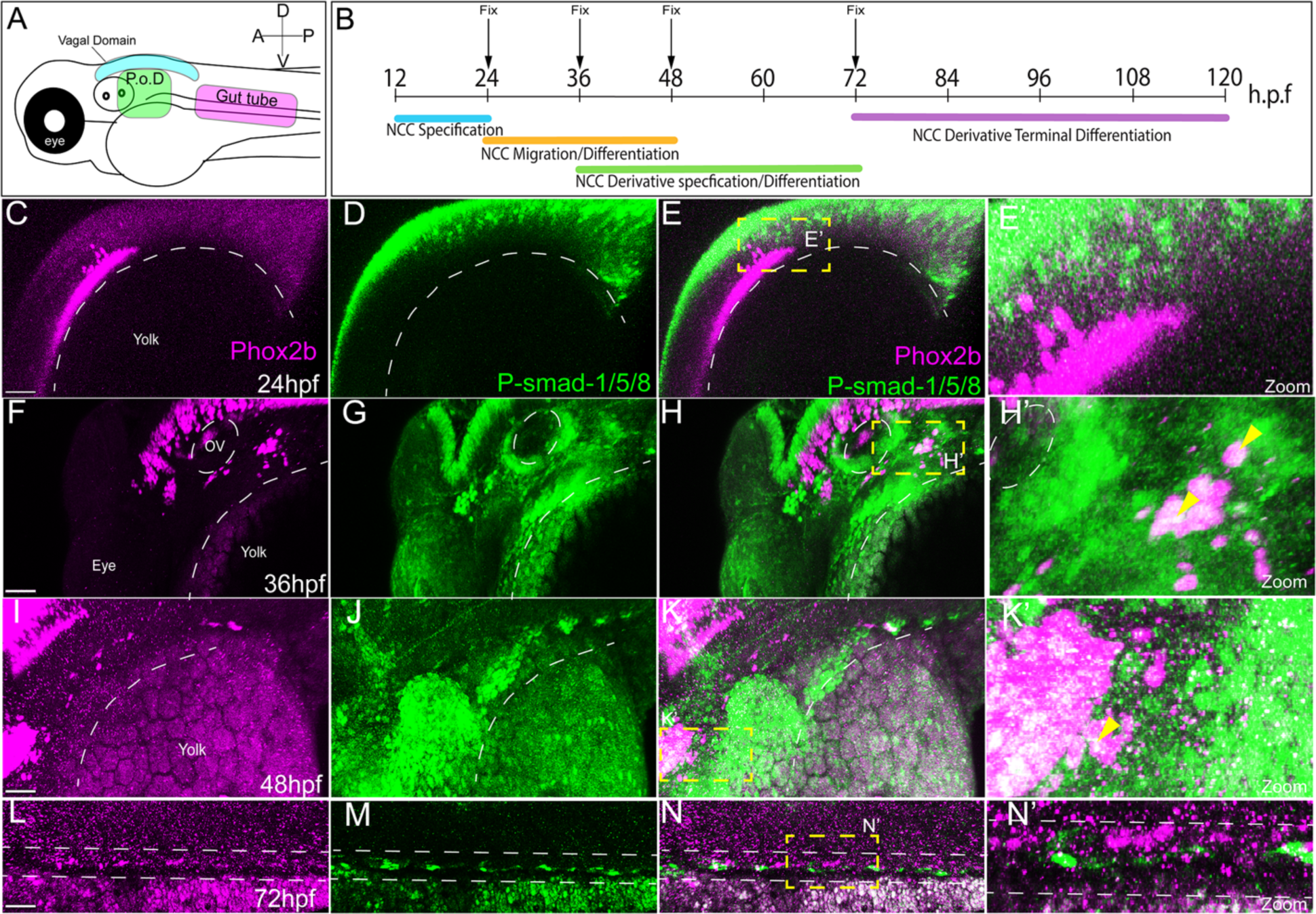
p-SMAD-1/5/8 is present in Phox2b^+^ enteric progenitors during early phases of enteric development. Schematic of regions of interest in the zebrafish embryo (A), and timepoints assayed for p-SMAD-1/5/8 and enteric neuronal differentiation markers (B). Immunohistochemical analysis of activated BMP signaling by immunostaining of p-Smad-1/5/8 and neurogenesis marker Phox2b at 24 (C-E’), 36 (F-H’), 48 (I-K’), and 72 hpf (L-N’’). Boxed region of co-expression of Phox2b and P-Smad1/5/8 (C’, F’,J’,M’). Anterior oriented to the left and dorsal to the top, in these images. Scale bar depicts 50 microns in C, F, I, L. P.o.D.: Post-otic Domain.

### Chemical attenuation of the BMP pathway alters the number and differentiation of phox2bb^+^ enteric progenitors in a time-dependent manner

A requirement for BMP signaling during ENS development has previously been determined by broad inhibition, via over-expression of endogenous inhibitors of BMP ligands in *Gallus gallus* (Goldstein et al 2005), and by gut explant treatment with BMP4 in *Mus musculus* (Chalazonitis et al., 2008, 2004). To further investigate when BMP signaling is required during vertebrate ENS development, we asked if BMP signaling perturbation alters ENS formation in zebrafish. To answer this question, we utilized chemical inhibition of BMP signaling via treatment with the potent and selective BMP antagonist K02288 (Sanvitale et al., 2013). For our experiments, inhibition began at 24hpf (Fig.4B), which is post-specification of the NCC born along the neural plate border, to avoid phenotypes specifically related to the developmental necessity of BMPs for early NCC specification (Das and Crump, 2012; Shih et al., 2017). When BMP signaling was attenuated between 24-120 hpf (Fig 4.B) there was a significant reduction in the number of −8.3*phoxbb*:Kaede^+^ (Kaede^+^) cells along the gut length, where larvae exhibited colonic aganglionosis, when compared with DMSO controls (Fig. 4A,F,L). Despite decreased numbers, the K02288-treated enteric cells exhibited neuronal differentiation, as assayed by the co-expression of Elavl3/4 and Kaede (Fig.4G,H,L), when compared to DMSO-treated controls (Fig.4C-E,L). When inhibition began at 36 hpf, Kaede^+^ cell numbers were similar to control numbers, however, their differentiation state was altered (Fig.4I-K,L). Specifically, while the colonization of the gut tube by Kaede^+^ cells was not significantly different than control, the fraction of Kaede^+^ cells expressing Elavl3/4 was significantly reduced (Fig.4L), indicating their failure to undergo neuronal differentiation. Overall, these results suggest a time-dependent necessity for BMP signaling during the orchestration of enteric progenitor numbers and/or neuronal differentiation along the gut.

**Figure 4.**
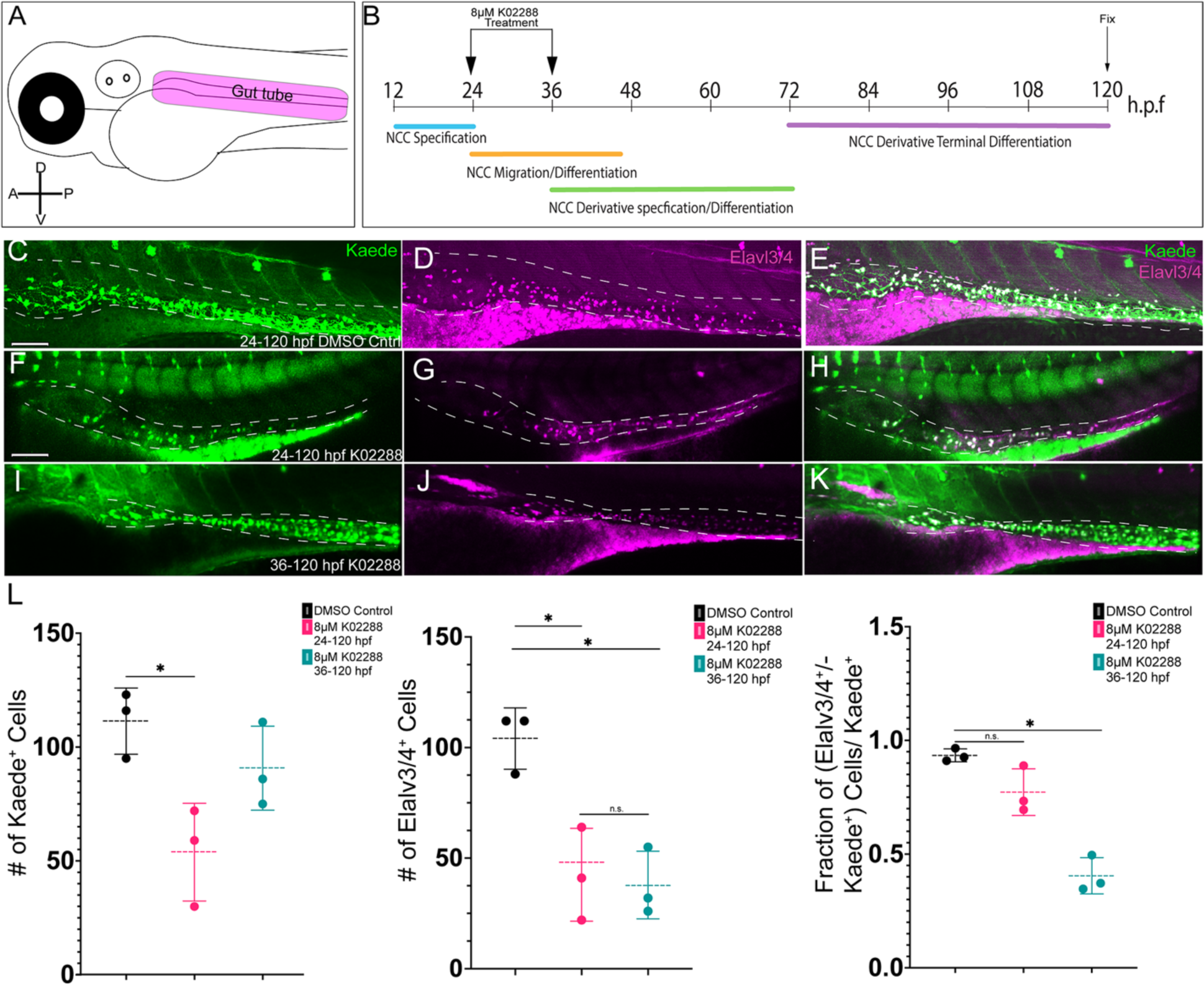
K02288 mediated attenuation of BMP signaling reduced enteric progenitor number in gut tube in a time-dependent manner. Schematic of the region of interest to be imaged for counting enteric neurons in the zebrafish embryo (A). Schematic of treatment paradigm with 8μM K02288 (B). Confocal imaging of the combinatorial expression of Kaede^+^ and Elavl3/4^+^ cells after 24-120 hpf DMSO control (C-E), or 24-120hpf (F-H), 36-120hpf (I-K) 8μm K02288 treatments. Counts of Kaede^+^ cells, Elavl3/4^+^ cells, or percentage of co-positive cells over Kaede^+^ cells using Imaris image analysis software (L). Anterior oriented to the left and dorsal to the top, in these images. Scale bar in C, F, I denotes 50 microns. * Denotes p-value < .05.

### bmp5, encoding for ligand BMP5, is a novel transcript identified and expressed in Phox2b^+^ enteric progenitors during zebrafish ENS development

Although BMP signaling has previously been identified as important for ENS development across organisms (Chalazonitis et al., 2008, 2004; Goldstein et al., 2005), and our current data demonstrate a temporal necessity for BMP signaling during zebrafish ENS development (Fig.4), investigations into which specific BMP ligand(s) are required for ENS development have been meager to date. Notably, Chalazonitis et al. have demonstrated, in *ex vivo* analysis, that treatment of NCC with BMP4 can alter the neuronal subtyping of subsequent enteric neurons. Following this, little has been done to identify additional novel ligands that may functionally play a role in development of the ENS.

To identify which BMP ligands were expressed during ENS development, we queried our single-cell *sox10*:GFP^+^ dataset, and specifically examined neural crest, enteric neural progenitors, and enteric neuron clusters (Fig.1). We identified the expression of *bmp5,* which encodes for a novel ligand throughout the enteric progenitor and neuronal populations (Fig.1C,E). Utilization of whole mount immuno-coupled hybridization chain reaction (WICHR) (Ibarra-García-Padilla et al., 2021) revealed strong expression of *bmp5* in the dorsal neural tube at 24 hpf, and in partial overlap with the expression domain of Phox2b^+^/Elavl3/4^+^ hindbrain (Fig.5A-D’). We detected co-expression of *bmp5* and Phox2b in enteric progenitors localized along the foregut during early enteric phases, 36-48 hpf, (Fig.5E-L’). Additionally, we identified expression domains of *bmp5* near the foregut at 36 hpf (Fig.5G). The expression of *bmp5* in enteric migratory routes, and enteric progenitors that migrate along the developing gut suggests possible functional roles for *bmp5* therein.

**Figure 5.**
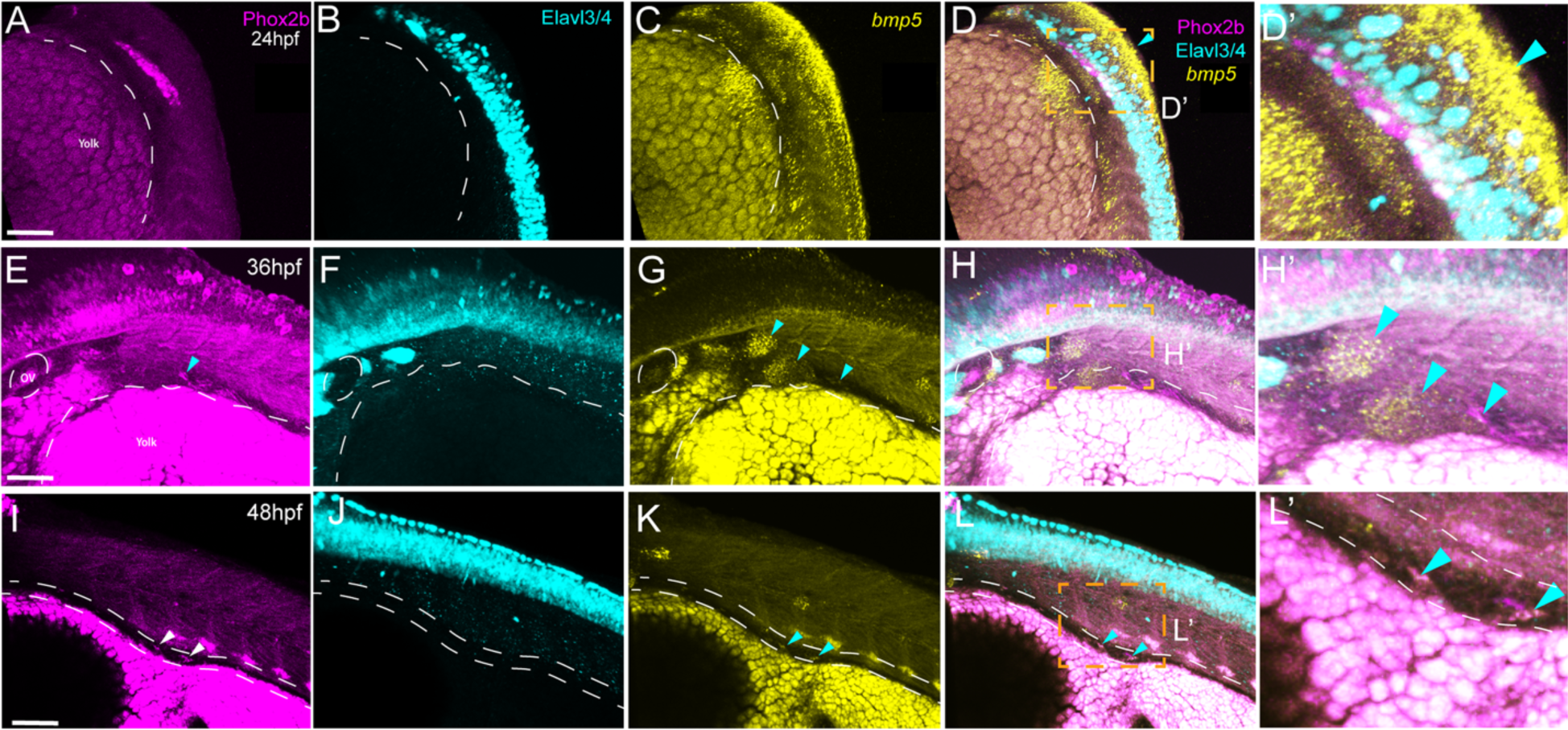
*bmp5* is expressed in and around the developing enteric progenitors during ENS development. Whole-mount immuno-coupled hybridization chain reaction analysis of novel ENS-related BMP ligand transcript, *bmp5*, in combination with neuronal markers Phox2b and Elavl3/4 at 24 hpf (A-D’), 36 hpf (E-H’), 48 hpf (I-L’). Co-expression of *bmp5* and enteric neurogenesis marker, Phox2b, is demonstrated by 48 hpf along the developing gut (L’). Anterior oriented to the left and dorsal to the top, in these images. Scale bar in A, E, I denotes 70 microns.

### Mutation in bmp5 reduced the number of −8.3phox2bb:Kaede^+^ cells along the zebrafish gut

Since *bmp5* is expressed around and within the developing ENS (Fig. 5), this suggests that it may play functional roles in the control of zebrafish ENS development. To elucidate if disruption in the *bmp5* genetic sequence leads to aberrations in ENS formation we utilized CRISPR-Cas9 mediated mutagenesis. To this end, a single guide RNA (sgRNA) was generated targeting the exon 1 of *bmp5* (Fig.6A) and injected with Cas9 into −8.3*phox2bb*:Kaede embryos. Injected embryos with indels were identified via PCR and T7 endonuclease activity (Fig.6B). To isolate mutations F0 −8.*3phox2bb*:Kaede adults were out-crossed to AB WT to generate F1 families. A subset of F1 embryos were dissociated for genomic DNA (gDNA) isolation and sequencing, which identified a 16 bp insertion within the *bmp5* locus, hereafter referred to as *bmp5^wmr2^* (Fig 6E). Examination of the ENS in *bmp5^wmr2/+^* larval fish revealed varied enteric phenotypes; while 80% of larvae were phenotypically WT, 20% of the larvae presented with enteric ganglionic phenotypes, where 16% presented with hypoganglionosis and 4% presented total aganglionosis (Fig.6C). Of particular interest, for larvae with hypoganglionosis, we observed a reduction in the number of *-8.3phox2bb*:Kaede^+^ cells along the length of the developing gut tube at 120 hpf, when compared with phenotypically WT siblings (*bmp5^+/+^*) (Fig.6D,E).

**Figure 6.**
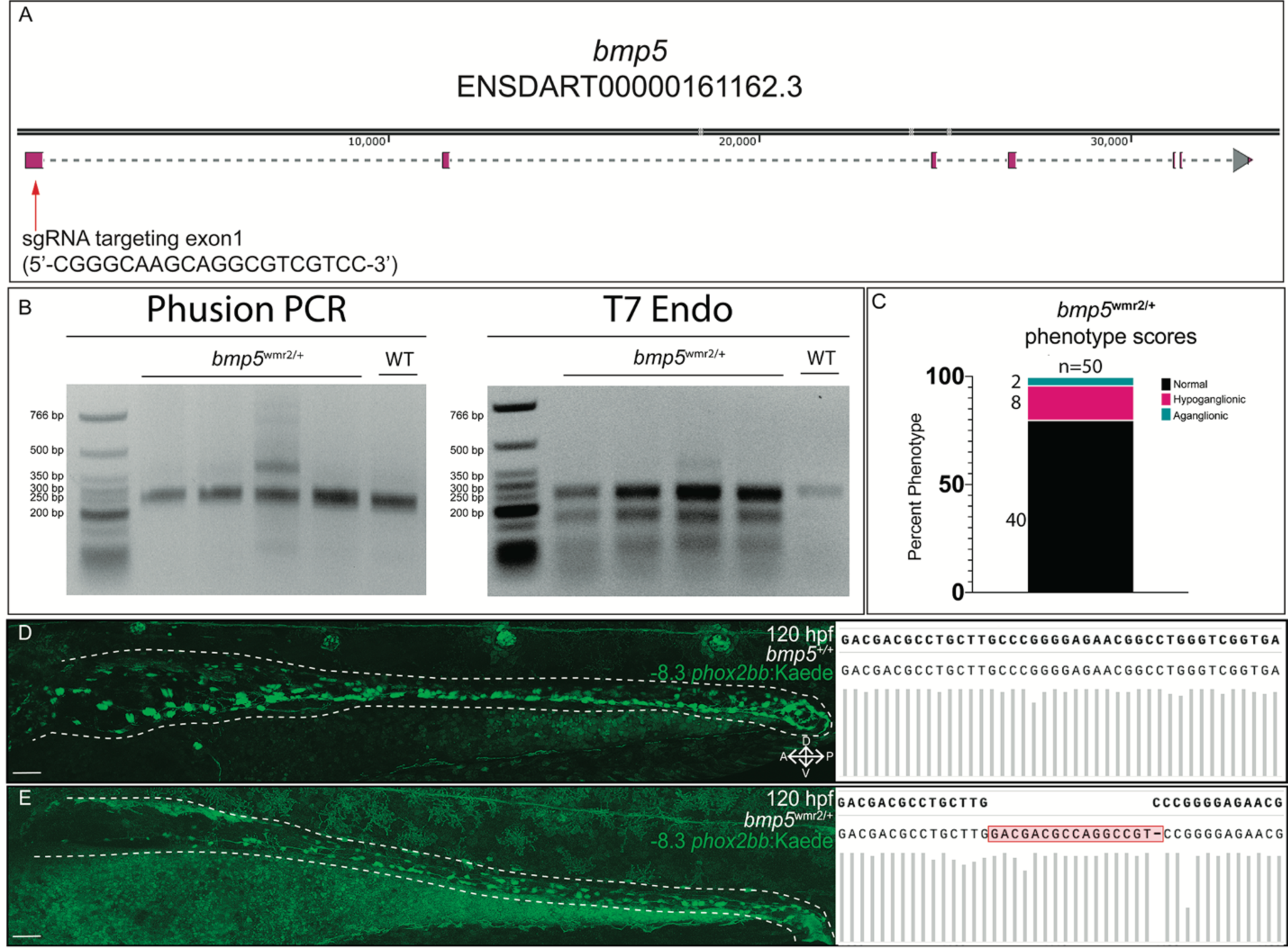
CRISPR mediated mutation in *bmp5* reduces *-8.3phox2bb*:Kaede^+^ enteric progenitors. Schematic of *bmp5* and CRISPR target region (A). Agarose gel analysis of *bmp5* target region amplicon via PCR and T7 endonuclease 1 activity (B). Fraction of ENS phenotypic distributions for *bmp5^wmr2/+^* larvae (C). Kaede^+^ cell reduction in *bmp5^wmr2/+^* when comparing *bmp5^+/+^* (D) to *bmp5^wmr2/+^* ENS (E) at 120 hpf, and the respective sequences around the targeted region of interest, with *bmp5^wmr2/+^* containing a 16-bp insertion. Anterior oriented to the left and dorsal to the top, in these images.

The reduction in enteric cell numbers in *bmp5* mutants, when compared to controls, suggests a requirement for proper *bmp5* function for normal colonization of the zebrafish gut. This may suggest a necessity for appropriate BMP5 function to properly and fully populate the length of the gastrointestinal tract. The successful generation of these novel mutants will allow for better understanding of BMP5 effects on later ENS development.

## Discussion

We have demonstrated in this study a novel role of Bone Morphogenetic Protein signaling in the context of the developing zebrafish enteric nervous system (ENS). We identified a time-dependent nature via which broad attenuation depletes progenitor number and/or neuronal differentiation of the cellular populations that give rise to the ENS. We also identified active BMP signaling, expression of BMP target genes, and for the first-time expression of, *bmp5*, encoding a novel BMP ligand in the ENS in and around the migratory enteric progenitors during early stages of ENS development. More specifically, we identified that a hypoganglionic gut is induced by generation of novel mutants in *bmp5^wmr2^*^/+^, thereby generating novel targets for the study of the development of the ENS.

Previous studies of BMPs in the development of amniote ENS have demonstrated that broad attenuation, via noggin overexpression, depletes HNK-1^+^ ENCC in the distal colon (Goldstein et al., 2005). Furthermore, targeted loss of *bmp2b* in zebrafish fully depletes the enteric neuron population (Huang et al., 2019), thereby suggesting functional roles for BMP *in vivo*. However, these roles have traditionally been relegated to the migratory nature of the pool of vagal neural crest that gives rise to the ENS. Due to the well-established timelines of enteric progenitor colonization of the gut (John B. Furness, 2006; Goldstein et al., 2005; Rao and Gershon, 2018; Young and Newgreen, 2001), our study identified windows in which BMPs effect could alter the proliferative capacity, specification state, or differentiation ability of enteric progenitors within the gut.

Specifically, our work here identified that chemical attenuation of BMP signaling in zebrafish starting at 24 hpf depleted enteric *-8.3phox2bb*:Kaede^+^ cells, however, these cells later differentiated into neurons (Fig.4). In contrast, with later BMP signaling reduction starting at 36 hpf, larvae appeared to have proper *-8.3phox2bb*:Kaede^+^ progenitor numbers, but many of those failed to differentiate into neurons by 120 hpf. These results suggest a role in which BMP early on, 24 hpf, is necessary for progenitor numbers, and later, 36 hpf, is needed for proper timing of enteric differentiation. This is supported by our expression analysis which identified dorsal neural tube expression of *bmp5* at 24 hpf (Fig.5), and by previous work demonstrating BMP5 necessity for early neural crest proliferation and survival (Shih et al., 2017). However, while we observed that the *bmp5* neural tube expression was lost by 36 and 48 hpf, it was expressed within both the tissue surrounding and within the Phox2b^+^ enteric progenitors, suggesting a potential role for *bmp5* in the timing of the differentiation of ENCCs, as demonstrated by our chemical attenuation. Combined, this temporal restriction of BMP signaling and *bmp5* spatiotemporal expression patterns suggest previously unknown ways BMPs, and in particular *bmp5,* may drive changes in ENS development and colonization by time-dependent control of either progenitor number maintenance, or timing of differentiation and/or specification of the progenitors.

Recent work identified *bmp2b* loss as a major contributor to the loss of migratory capacity of ENCCs into the zebrafish developing gut tube (Huang et al., 2019). However, this loss in particular could be associated with general migratory loss of the neural crest populations (Correia et al., 2007) and not specifically loss of ENCC migration. It is known that BMPs play varied roles throughout development depending upon the context, time, and binding partners (Correia et al., 2007; Das and Crump, 2012; Pomreinke et al., 2017; Zuzarte-Luís et al., 2004). Previous transcriptomic (Memic et al., 2018) and *in vivo* (Shih et al., 2017; Wu et al., 2014) analyses have identified *BMP5* expression in the lineages of cells that give rise to the ENS or in tissues where enteric progenitors will migrate. These data support the hypothesized expression of BMP5 in the developing human and mouse gut (Memic et al., 2018). Our study identified novel expression of *bmp5,* and many BMP target genes, in and around the migratory ENCCs during active ENS colonization, and our novel zebrafish *bmp5*^wmr2/+^ mutants demonstrate alterations in the developing ENS cell numbers.

Collectively this study, with previous work by others, supports a temporally dependent role for broad BMP activity, and novel role for BMP5 in regulating progenitor numbers during development of the ENS. Combined with our strong transcriptomic, *in situ* and *in vivo* analysis we have additionally identified novel candidates that may help orchestrate the complex balance of proliferation, differentiation, and migration necessary for proper ENS colonization.

## Materials and Methods

### Zebrafish husbandry, and embryo and larvae collection

This work was conducted in accordance with the Institutional Animal Care and Use Committee (IACUC) of Rice University. Embryos and larvae for all experiments were collected from controlled breeding of adult zebrafish for synchronous staging. All embryos were maintained at 28°C in standard E3 embryo medium until 24 h post fertilization (hpf), then were transferred to 0.003% 1-phenyl 2-thiourea (PTU)/E3 solution (Karlsson et al., 2001), Transgenic embryos used for this work include *−8.3phox2bb*:Kaede^em2Tg^ (Harrison et al., 2014) and *−8.3phox2bb*:Kaede;*bmp5^wmr2/+^* fish (this study). Embryos and larvae were collected out of their chorions at the stage noted in each experiment.

### Transcriptomic expression analysis

Neural crest, enteric progenitors, and enteric neuron cell type identities based on marker gene transcript expression in zebrafish have previously been described in *sox10*:GFP single-cell RNA-seq atlases (Gene Expression Omnibus (GEO) database accession number: GSE152906 and GSE163907) (Howard et al. 2021; Lencer, Prekeris, and Artinger, 2021). Using Seurat as previously described (Howard et al. 2021; Butler et al. 2018), subsetting for the identity populations was performed to assay target gene expression at 24, 48, 68 hpf during early, mid and late ENS development, respectively, where we then queried and identified 37 BMP pathway associated genes (zfin.org) expressed. Data was visualized in subsetted feature plots, or dot plots, as previously described (Howard et al. 2021).

### CRISPR-Cas9 guide RNA design and synthesis

A sgRNA targeting exon 1 of *bmp5* (5’-CGGGCAAGCAGGCGTCGTCC -3’) was designed by manually searching the *bmp5* CDS (ENSDART00000161162.3) for protospacer adjacent motifs (PAM) within CDS domains. The generation of the *bmp5* sgRNA based on previously described work (Zhang et al., 2020). Briefly, the sgRNA was generated from a custom DNA oligo containing an SP6 promoter, as previously reported (Gagnon et al., 2014), purified and resuspended in nuclease-free water. Quality of sgRNA stock was assessed via agarose gel electrophoresis and nanodrop quantitation.

### CRISPR-Cas9 microinjection, *bmp5^wmr2/+^* fish line establishment and genotyping

Zebrafish embryos obtained from in-crossing Tg(−8.3*phox2bb*:Kaede)^+/−^ adults were injected at the one-cell stage with a cocktail containing: phenol red dye .5 nanoliter (nl), 966 picogram (pg) New England Biolabs Cas9 (M0646T) and 120 pg of sgRNA per embryo targeting exon 1 of *bmp5* (5’-CGGGCAAGCAGGCGTCGTCC -3’) (as described above). Injected F0 larvae were screened at 5 dpf for ENS defects in which regions of the intestine demonstrated reduced Kaede^+^ cells, and a subset of F0 embryos were dissociated and used in T7 endonuclease activity assays (NEB E3321) in order to validate successful CRISPR-Cas9 activity, as previously described (Baker et al., 2022). Injected F0 larvae exhibiting WT phenotype were collected and raised in our in-house zebrafish facility, and when reaching sexual maturity were outcrossed to AB WT. A subset of F1 larvae were dissociated and used in T7 endonuclease activity assays and/or imaged (Fig.6), and remaining F1 fish grown to adults were fin clipped and genotyped (Meeker et al., 2007) using forward (5’-GGACTTCTGTGGAGCTGTTTAG-3’) and reverse (5’-CCCCAGTATCTCGTTCTCCTCTT-3’) primers to produce a 257 base pair (bp) amplicon from the *bmp5* locus. To obtain sequence of mutation the previously described 257 bp amplicons were cloned into pCR-Blunt II-TOPO vector using the manufacturer’s instructions (Invitrogen, 45-0245). Zero Blunt TOPO reactions were transformed into DH5α bacteria. Bacteria were mini-prepped and screened using restriction digest cloning. Presumptive plasmids containing *bmp5^wmr2/+^* and *bmp5^+/+^* were confirmed following whole-plasmid sequencing by Plasmidsaurus. F1 heterozygous adults harboring 16 bp insertion at bp position 400 of the *bmp5* coding domain sequence (as described in results) in *bmp5* were designated *bmp5^wmr2/+^* fish. The 16 bp insertion resulted in the introduction of a predicted premature stop codon at bp position 521.

### Live Confocal microscopy and image exports

Larvae were anesthetized using 0.4% Tricaine (Sigma-Aldrich, A5040), then mounted in 15 µ-Slide 4 Well imaging chambers (Ibidi, 80427) using 1.0% low melt temperature agarose dissolved in E3 media. Embedded fish were then covered in 1× PTU/E3 media supplemented with 0.4% Tricaine. Confocal microscopy was performed using an Olympus FV3000 and FluoView software (2.4.1.198), using a long working distance 20.0× objective (UCPLFLN20X) at a constant temperature of 28°C using OKOLAB Uno-controller imaging incubator. Final images were combined in the FluoView software and exported for analysis IMARIS image analysis software (Bitplane). Figures were prepared in Adobe Photoshop and Illustrator software programs.

### Zebrafish treatment with Chemical inhibitor

K02288 (SML1307, Sigma Aldrich) (Sanvitale et al., 2013) master stock was diluted in DMSO and then further diluted in 1xPTU/E3 medium to give the required inhibitor concentration. Approximately 20 *−8.3phox2bb*:Kaede^+^ embryos per well, in a volume of 1 mL of 1xPTU/E3 medium supplemented with DMSO or 8 micromolar K02288, were placed in 12-well plates at 24 hpf or 36 hpf, and incubated until 120 hpf. Larvae were processed and imaged with confocal microscopy, as described above.

### Enteric cell counting

Cell counting was performed using spots function in Imaris (10.0.0). The position of every ENCC body (spot) was identified across *x*-*y*-*z* planes using manual spot detection in the Kaede, Alexa flour-488, or Alexa flour-647 channels to determine the number of Kaede^+^, Elavl3/4^+^, and Kaede/Elavl3/4 co-positive ENCCs (spots) at 120 hpf. Every ENCC was labeled using unbiased spot determination, then manually, spots not recognized within the gut we added along the A-P axis. Co-positive ENCCs (spots) were determined using the Imaris colocalization function. Total spots were exported from Imaris and further analyzed for graphical depictions using GraphPad Prism (version 9.5.1[528]).

### Hybridization chain reaction, Whole Mount Immunofluorescence, WIHCR

Hybridization chain reaction (HCR) and Whole Mount Immuno-Couple HCR (WIHCR) experiments were performed in accordance with previously described methods (Howard et al., 2021; Ibarra-García-Padilla et al., 2021) for the following transcripts with RefSeq IDs: *msx1b*, NM_131260.1; *id2a*, NM_201291; *crestin*, NM_130993; *hand2*, NM_131626; *bmp5*, NM_201051. Whole-mount immunofluorescence experiments were conducted according to the methods previously described (Uribe and Bronner, 2015). The following primary antibodies were used: mouse monoclonal IgG1 anti-Phox2b (B-11, Santa Cruz Biotechnology, SC-376997, 1:250), rabbit polyclonal anti-p-Smad1/5/9 IgG (Cellular Signaling Technologies, 13820S), rabbit polyclonal anti-Kaede IgG (MBL International, PM102M, 1:250), and mouse monoclonal anti-Elavl3/4 IgG2b (Invitrogen Molecular Probes, A21271, 1:250). The following secondary antibodies were used from Invitrogen: Alexa Fluor 488 goat anti-rabbit IgG (A11008, 1:500), Alexa Flour 488 goat anti-mouse IgG1 (A21121, 1:500), Alexa Fluor 568 goat anti-mouse IgG1b (A21124, 1:500), and Alexa Fluor 647 goat anti-mouse IgG2b (A21242, 1:500). Confocal microscopy was performed using an Olympus FV3000 and FluoView software (2.4.1.198), using a long working distance 20.0× objective (UCPLFLN20X). Final images were combined in the FluoView software and exported for analysis IMARIS image analysis software (Bitplane). Figures were prepared in Adobe Photoshop and Illustrator software programs.

### Statistics

Statistical analysis was performed in GraphPad Prism (version 9.5.1[528]). For comparisons, data was tested using two-tailed unpaired *t*-test, \**P*<0.05, n.s., non-significant (*P*<0.05).

## Acknowledgments

Our great thanks go to the Rice University Shared Equipment Authority and Dr. Alloysius Budi Utama for assistance, training, and use of the Imaris workstation for Imaris image analysis performed within this study. We thank Sage Simmons for their insight and advice. We also thank Tobechukwu Dubem Nwigwe and Jhené Aiko Efuru Chilombo for technical assistance.

---

**Figure Supplement 1.**
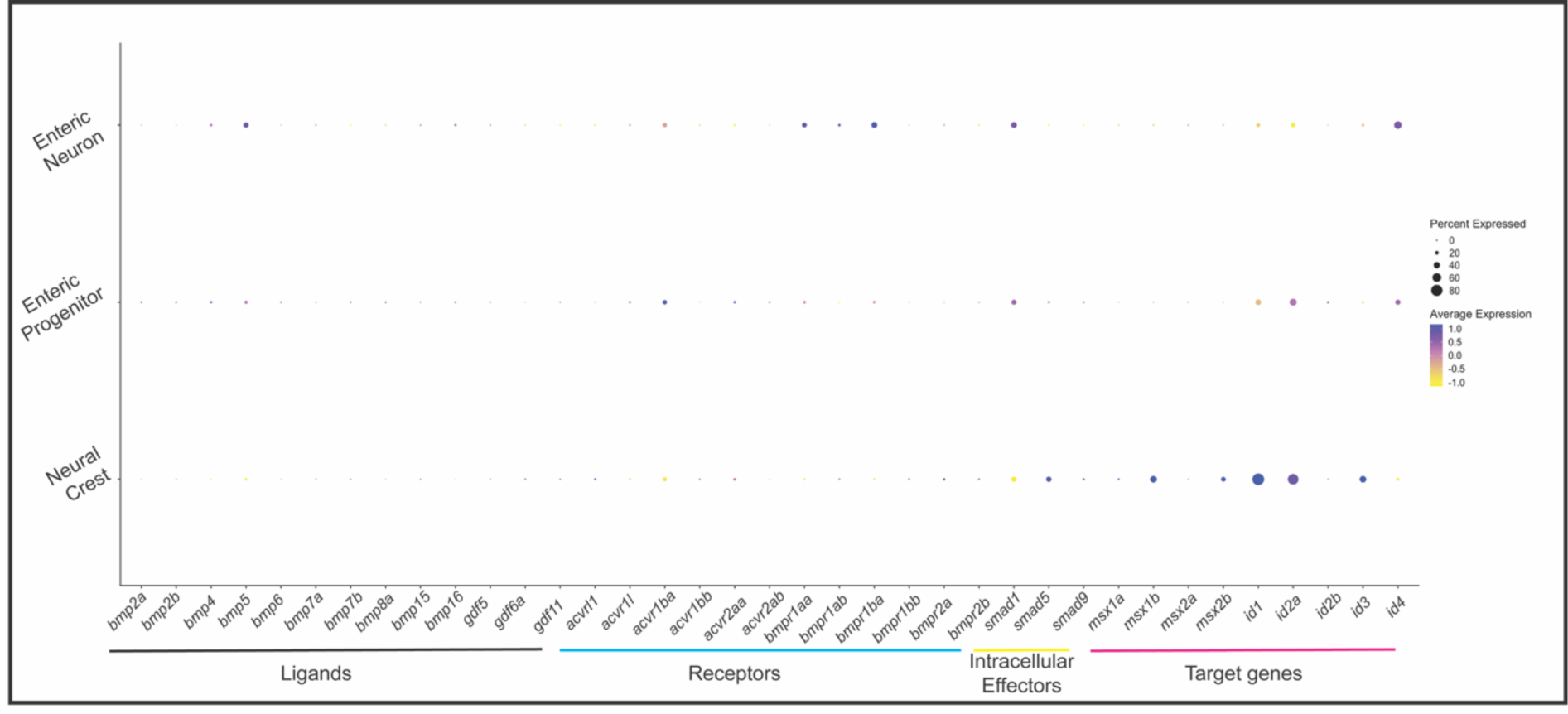
37 Bone Morphogenetic Signaling pathway members expressed across *sox10* derivatives. Dot plot depicts the transcript expression of BMP ligands, receptors, intracellular targets, and target genes across the neural crest, enteric progenitors, and enteric neurons, as queried from datasets from Howard et al. 2021.

